# *In silico* analysis of the evolution of root phenotypes during maize domestication in Neolithic soils of Tehuacán

**DOI:** 10.1101/2024.11.18.623787

**Authors:** Ivan Lopez-Valdivia, Miguel Vallebueno-Estrada, Harini Rangarajan, Kelly Swarts, Bruce F. Benz, Michael Blake, Jagdeep Singh Sidhu, Sergio Perez-Limon, Ruairidh J. H. Sawers, Hannah Schneider, Jonathan P. Lynch

**Author notes:** Corresponding author: Jonathan Lynch.

## Abstract

Roots are essential for plant adaptation to changing environments, yet the role of roots in crop domestication remains unclear. This study examines the evolution of root phenotypes from teosinte to maize, a transition resulting in reduced nodal root number (NRN), multiseriate cortical sclerenchyma (MCS), and increased seminal root number (SRN). We reconstructed the root phenotypes of maize and teosinte, as well as the environments of the Tehuacan Valley over the last 18,000 years using a combination of ancient DNA, paleobotany, and functional-structural modeling. Our models reveal that increasing Holocene atmospheric CO_2_ concentrations favored the appearance of reduced NRN and MCS between 12000 to 8000 years before present (yBP), promoting deeper root systems. The advent of irrigation by 6000 yBP switched nitrogen distribution from topsoil to subsoil domains, a change which increased the utility of reduced NRN and MCS. Comparison of allelic frequencies among ancient samples ranging from 5500 to 500 yBP suggest that increased SRN may have appeared around 3500 yBP, coinciding with a period of increased human population, agricultural intensification, and soil degradation. Our results suggest that root phenotypes that enhance plant performance under nitrogen stress were important for maize adaptation to changing agricultural practices in the Tehuacan Valley.

**Classification:** Physiology & Development

## Introduction

Maize (*Zea mays ssp. mays*) was domesticated from teosinte (*Zea mays ssp. parviglumis*) in central Mexico around 9,000 years before present (yBP) (Matsuoka *et al*., 2002; Yang *et al*., 2023). While the importance of roots for environmental adaptation is well-established (Lynch *et al.,* 2022; McLaughlin *et al.,* 2024), their specific role in maize domestication remains poorly understood (Schmidt et al., 2016). Compared with teosinte, maize exhibits distinct root phenotypes, including greater seminal root number (SRN), reduced nodal root number (NRN), and the development of multiseriate cortical sclerenchyma (MCS) in nodal roots (Burton *et al*., 2021, Schneider *et al*., 2021). However, the functional utility of SRN, NRN, and MCS in the context of maize domestication remains unclear.

The domestication of maize coincided with the end of the Pleistocene, a period marked by rising atmospheric temperatures and CO_2_ concentrations (Piperno *et al*., 2007; Ahn *et al*., 2004; Correa-Metrio *et al*., 2012). Experiments in controlled environments showed that teosinte grown under late Pleistocene and early Holocene atmospheres exhibited maize-like vegetative and reproductive phenotypes that presumably improved the harvesting effectiveness of teosinte, making it attractive to early cultivators (Piperno *et al*., 2015; Piperno *et al*., 2019). Eventually, those phenotypes were likely fixed by genetic assimilation in modern maize populations (Lorant *et al*., 2017). Although elevated CO_2_ is known to promote root growth in maize (Hiltpold *et al*., 2020), the impact of these historical CO_2_ fluctuations on root phenotypes of domesticated crops is unknown.

The emergence and intensification of agriculture in the Holocene also changed the soil environment. Soil cultivation without concomitant use of organic soil amendments degrades soil fertility by accelerating microbial oxidation of soil organic matter (*Sandor et al*., 1986a,b). Soil degradation associated with early Holocene agriculture is evident (McAuliffe *et al*., 2001), as is decreased plant nitrogen status following crop domestication (Sandor and Gersper, 1988; Araus *et al*., 2014). Furthermore, irrigation intensifies soil erosion as well as nitrogen leaching (Graham *et al*., 2022). The importance of these processes in early maize domestication environments is shown by the substantial soil erosion (McAuliffe *et al*., 2001; Cook, 1949), and archeological evidence of ancient irrigation systems (Dillehay *et al*., 2005; Neely *et al*., 2022; Borejsza *et al*., 2014; Cajigas *et al*., 2020; Huckleberry *et al*., 2014; Doolittle *et al*., 2006). The use of ancient irrigation systems would relieve plants of water stress but would decrease soil fertility due to soil erosion and nitrogen leaching.

These soil and atmospheric changes likely influenced the evolution of maize root systems. For example, increased SRN, which relies on seed reserves, improves shoot growth under suboptimal nitrogen and phosphorus availability (Perkins and Lynch, 2021). Maize genotypes with reduced NRN decreased interplant competition and improved nitrogen capture at high plant densities (York *et al*., 2015; Gaudin *et al.,* 2011). Maize lines with MCS have greater root lignification, which improves penetration ability, promoting greater root depth and greater biomass in compacted soils (Schneider *et al*., 2021). Interestingly, 5000-year-old maize roots from the Tehuacan Valley show MCS and reduced NRN, but not increased SRN (Lopez-Valdivia *et al*., 2022). This finding raises questions about the evolutionary sequence of root phenotypes and the environmental pressures that drove these changes. The Tehuacan Valley offers a unique window into this process, with its extensive archaeological record documenting the transition from hunter-gathering to farming over the last 10,000 years (MacNeish, 1967). This transition is described by a gradual increase of consumption of ancestral crops species, followed by the intensification of agriculture by the construction of ancient irrigation systems (Neely *et al., 2022*) and the consequent soil degradation (McAuliffe *et al.,* 2001). Despite this detailed archeological record, it remains unclear why reduced NRN and MCS appeared earlier than increased SRN, and what specific environmental factors drove their emergence.

Atmospheric and soil dynamics may have influenced the fitness landscape of root phenotypes, promoting the appearance of SRN, NRN, and MCS in modern maize. *In silico* approaches to this problem are needed since Holocene agricultural environments and transitional phenotypes no longer exist. *OpenSimRoot* is a functional-structural plant/soil model that simulates root growth and soil resource capture by whole plants interacting dynamically with soil and atmospheric environments (Postma *et al*., 2017). *OpenSimRoot* simulates plant and soil responses to drought, nutrient availability, mechanical impedance, atmospheric CO_2_ and temperature, including dynamic interactions between local and global plant responses to root water capture, soil water status, and soil mechanical impedance, making it a suitable tool in this context (Schäfer *et al*., 2022; Strock *et al*., 2022; Ajmera *et al*., 2022).

We hypothesize that root phenotypes adapted to anthropogenic changes in the soil environment caused by the advent of agriculture in the Holocene (Figure 1). In this study, we used ancient DNA, paleobotany, and functional-structural modeling to estimate the approximate time of appearance of modern SRN, NRN, and MCS phenotypes. We reconstructed soil and atmosphere environments of the Tehuacan Valley for the last 18,000 years using archeological and paleoclimatic data. We used *OpenSimRoot* and ancient DNA to simulate the growth of teosinte, maize, and transitional phenotypes under Holocene environments and evaluate their performance. Finally, we dissected the phenotypes and the environments to determine what elements of the phenotype/environment interaction were most important in shaping plant adaptation.

**Figure 1.**
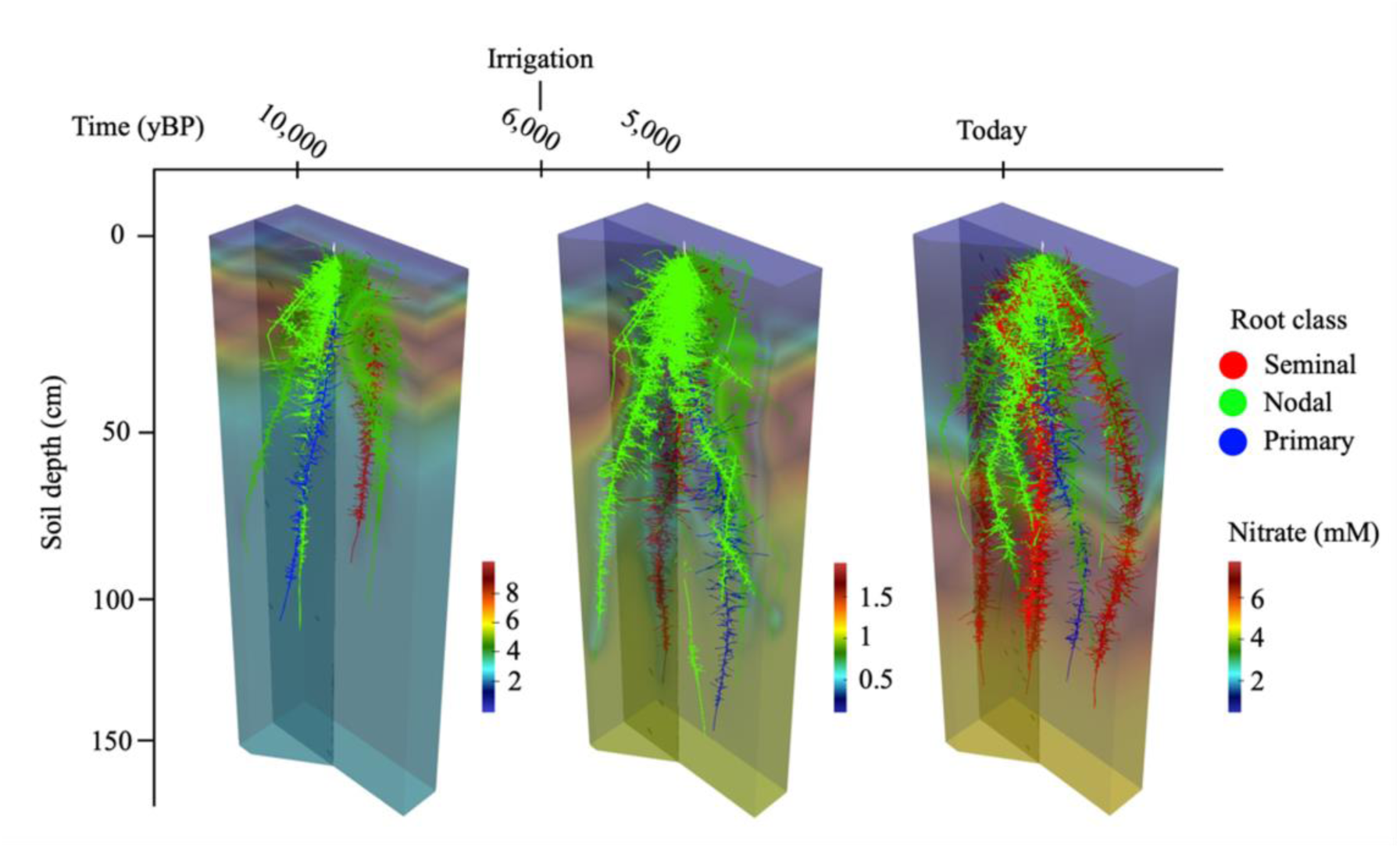
Evolution of root phenotypic transition from teosinte to maize in the last 10,000 years. Simulated root phenotypes using the functional-structural model *OpenSimRoot*. The root phenotype of 5,000 yBP includes MCS, which increases root depth. Nitrogen depletion by leaching and root capture is shown as a gradient. Nitrogen concentration is shown in mmol of nitrate per l of soil solution. Dimensions of the soil matrix are 150 x 50 x 25 cm. Irrigation started by 6,000 yBP. Simulation of 40 days of growth. A video animation is provided in Supplementary Movie 1.

## Materials and Methods

### Root phenotype parameterization

#### Seminal root number and nodal root number

The phenotypes that distinguish modern maize root system from teosinte are greater seminal root number (SRN), smaller nodal root number (NRN), and the presence of multiseriate cortical sclerenchyma (MCS). We used *OpenSimRoot* to simulate the root phenotypes of maize, teosinte, and ancient transitional phenotypes represented as the combination of teosinte phenotypes plus modern maize phenotypes, for instance: teosinte+SRN, teosinte+NRN, teosinte+MCS, teosinte+SRN+MCS, teosinte+NRN+SRN and teosinte+NRN+MCS. To simulate seminal root number, we considered the mean SRN for a diverse population of maize landraces and teosintes; 4 seminal roots for maize and 1 seminal root for teosinte (Burton *et al*., 2013). Maize populations have 4 fewer nodal roots than teosinte populations at stage V6-V7 (Burton *et al*., 2013). We simulated 20 and 24 nodal roots for the first 5 nodes for maize and teosinte, respectively.

#### Multiseriate cortical sclerenchyma

To simulate MCS we measured the carbon density in MCS and non-MCS maize roots. Twelve IBM (intermated B73*MO17) recombinant inbred lines were grown at the Russell E. Larson Agricultural Research Farm at Rock Springs, PA, USA (40°42’40.915” N, 77°,57’11.120” W) from June through September 2021. We employed a randomized complete block design with four blocks. Within each block, all the 12 genotypes were randomized. Before planting, all fields were fertilized to meet the nutrient requirements of maize as determined by soil tests at the beginning of the season. Each genotype was planted in a three-row plot with a length of 4.6 m long and 76 cm row spacing at a density of 73,300 plants ha^-1^. As needed, irrigation was supplied throughout the season. At approx. 80 days after planting (post-anthesis), one plant per plot was harvested following the shovelomics protocol (Trachsel *et al*., 2011). Root anatomy samples of the second and fourth node roots (3 cm away from the base) were collected. Anatomy samples were immediately stored in 75% ethanol and later processed using laser ablation tomography (LAT) (Strock *et al*., 2019). Samples were divided into MCS or no-MCS categories (Schneider *et al*., 2021). In total, 11 non-MCS and 19 MCS root samples were used. One cm long segment for each sample was saved and dried at 60 C for 24 hours. The dried samples were weighed using PerkinElmer Ultra micro balance AD-6 (C099-1141). Tissue density was calculated by dividing tissue dry weight by tissue volume.

A greenhouse mesocosm experiment was conducted to measure root respiration in optimal and water deficit conditions. Six IBM recombinant inbred lines (IBM146, IBM14, IBM86, IBM284, IBM178, IBM284) were grown from April 14 to May 12 2020 in four replications in a randomized complete block design. The greenhouse was located on the campus of Pennsylvania State University in University Park (40° 489 N, 77° 519 W) and plants were grown under constant conditions (14 h of day at 28°C/10 h of night at 24°C, 40%–70% relative humidity). Two seeds were transplanted into individual mesocosms consisting of polyvinyl chloride cylinders with an inner diameter of 15.5 cm and height of 1.54 m and lined with transparent 6 mm high-density polyethylene film to aid with root sampling. Each mesocosm was filled with a 30 L mixture consisting of 50% commercial grade medium sand (Quikrete), 27% horticultural grade fine vermiculite (D3, Whittemore Companies Inc.), 18% field soil (Hagerstown silt loam: fine, mixed, semi-active, mesic Typic Hapludalf, air-dried, crushed, and sieved through a 4-mm mesh), and 5% horticultural grade super-coarse perlite (Whittemore Companies Inc.), by volume. Mineral nutrients were provided by mixing the medium with 70 g per column of Osmocote Plus fertilizer consisting of 15% (w/w) N, 9% (w/w) P, 12% (w/w) potassium, 2.3% (w/w) sulfur, 0.02% (w/w) boron, 0.05% (w/w) copper, 0.68% (w/w) iron, 0.06% (w/w) manganese, 0.02% (w/w) molybdenum, and 0.05% (w/w) zinc (ScottsSierra Horticultural Products). A thin surface layer (∼1 cm) of perlite was added to each mesocosm to help retain moisture and reduce compression of media. Two days before planting, each mesocosm was irrigated with 4.5 L of tap water. In the first 4 d, plants received 100 mL of deionized water every day. Then, 200 mL of deionized water was irrigated for the control treatment every 2 d, and the water deficit treatment received no further irrigation. After one week, plants were thinned in individual mesocosms to one seedling.

At harvest, the plastic liner was removed from the PVC mesocosm and cut open, and roots were separated from the soil by vigorous rinsing at low pressure with water. Root respiration of axial roots was measured. Three representative 10-cm root segments from roots emerging from the fourth node were excised 5-8 cm from the base. Excised root samples were patted dry and placed in a 40-mL custom chamber connected to the LI-6400 infrared gas analyzer (Li-Cor Biosciences) separately. The temperature of the chamber was maintained at 26°C using a water bath while respiration was measured. Carbon dioxide evolution from the root segments was recorded every 5 s for 180 s. Afterwards, root samples were preserved in 70% EtOH for laser ablation tomography and MCS imaging and phenotyping (as described above). For MCS we modeled a 22% greater penetration ability in axial roots (Schneider *et al*., 2021) represented as a reduction of the elongation rate at 4000 kPa mechanical impedance with MCS and 1000 kPa for non-MCS phenotypes as described by (Strock *et al*., 2022).

#### Kernel weight model

Kernel weight is a phenotype that changed during domestication and is a key factor determining the utility of SRN (Perkins *et al*., 2021). To determine the effect of kernel weight on the selection of modern root phenotypes during the Holocene we constructed a statistical model to estimate kernel weight over time based on the glume width and depth of paleobotanical maize cobs of different ages. To build the model we measured the weight of 50 seeds of 143 families of the Multiparent Advanced Generation Inter-Cross population (MAGIC) created from 8 native maize varieties from Mexico. We measured 3 cobs per family. In every cob, we measured the length and width of 3 glumes. Glumes were selected from the apical, central, and basal regions. The average of the 3 glumes along the 3 sections of the 3 cobs was considered a representative of the glume size per family (Data S1). We measured the glume width and length of ancient cobs Pur26, Cox35, Cox31, Cox14, Pur78, Pur19, SM6, SM8, SM7, SM9, SM10, Tehuacan162 from Tehuacan Valley and Naquitz_D10, Naquitz_C9 from Guila Naquitz utilizing images available in (Torres-Rodríguez *et al*., 2018; Ramos-Madrigal *et al*., 2016) and (Piperno *et al*., 2001), respectively. We trained and tested a multiple linear regression model in R version 4.2.3 (R Core Team, 2019) utilizing the glume width and length of the MAGIC population to predict seed weight (RMSE = 0.029 g). We used the model to predict the seed weight of the ancient cobs based on the glume size (Data S2).

### Environmental parameterization

To reconstruct the soil and atmospheric environment of the late Pleistocene and Holocene periods in the Tehuacan Valley (18°27′42″N, 97°23′34″W) we integrated climatic and paleoclimatic data into *OpenSimRoot*. To simulate the representative atmospheric patterns of a regular growing season (June 19 – August 18) we obtained the precipitation, minimum temperature, maximum temperature, wind speed, and relative humidity from 1980 to 2021 (Data S3) from POWER Data Access Viewer v2.0.0 (https://power.larc.nasa.gov/data-access-viewer/ consulted on 2023/3/17) and used the average as the representative atmosphere (Data S4). To simulate the atmosphere of the last 18,000 yBP we considered intervals of 2,000 years and adjusted the average patterns of the Tehuacan Valley based on the paleoclimatic events of precipitation (Hodell *et al*., 2008), temperature (Correa-Metrio *et al*., 2012), and atmospheric CO_2_ (Ahn *et al*., 2004). To simulate soil texture we used core data from SoilGrids (Poggio *et al*., 2021), core coordinates -97.3340, 18.3759, core number 215588, MX-INEGI dataset. We calculated the van Genuchten parameters (Data S5) utilizing Rosetta3 (https://www.handbook60.org/rosetta/). We simulated irrigation events starting at 6,000 yBP (Neely *et al*., 2022) and lasting until the present by increasing the precipitation patterns by a factor of 1.25 for 6,000 yBP, 1.5 for 4,000-2,000 yBP, and 2 for the modern environment (Data S6). We represented soil degradation as a 50% decrease in organic matter and 90% decrease in mineralization (Sandor *et al*., 1986b; Sandor and Gersper, 1988). Today’s environment was simulated using the actual environmental conditions of Tehuacan Valley with and without irrigation (Figure S4). We used today’s environment with irrigation for the simulations across the article. The future scenario was simulated based on CO_2_, temperature, and precipitation projected in the Sixth Assessment Report from the Intergovernmental Panel on Climate Change (Lee *et al*., 2021). The compiled phenotypic, atmospheric, and soil data to parameterize the model with transitional root phenotypes and Tehuacan Valley environment across time is available in the following repository: https://github.com/ilovaldivia/Lopez-Valdivia_2023_RootEvolution.git.

### OpenSimRoot runs

To simulate the root growth of teosinte, maize, and transitional domesticates in late Pleistocene and Holocene environments, we used the functional-structural model *OpenSimRoot* available at https://rootmodels.gitlab.io/. We used the OpenSimRoot_v2 (Schäfer *et al*., 2022) with git version 9621150b6f due to its new capabilities to simulate drought responses. We ran the simulations utilizing High-Performance Computing at Pennsylvania State University. All the input files required to replicate our results are available in the following repository: https://github.com/ilovaldivia/Lopez-Valdivia_2023_RootEvolution.git. A detailed description of *OpenSimRoot* modules can be found in Postma *et al*., (2017) and a manual on how to run simulations can be found in Schäfer *et al*., (2022).

### Extraction UDG library build and sequencing of ancient samples

Sample processing and DNA extraction were performed following all necessary procedures to avoid human-related or cross-sample contamination in a clean laboratory optimized for paleogenomics, as previously described (Vallebueno-Estrada *et al*., 2013; Vallebueno-Estrada *et al*., 2023). Archeological maize specimens (Data S7) were processed at the ancient DNA facilities of Vienna University, Austria. DNA extraction was 15 mg of the inner parenchyma tissues. Isolation of DNA was carried out in clean laboratory facilities with dedicated reagents and equipment that are frequently sterilized, and UV treated. To prevent cross-sample and human-related contamination, we used new disposable plastic material and filtered pipette tips, and personnel wear protective gear such as full bodysuits, masks, and doubled gloves. Work was conducted in laminar flow hoods. Samples were ground with a mortar and pestle. DNA extraction was conducted using a freshly prepared PTB lysis buffer (PTB 2.5mM, DTT 50mM, Proteinase K 0.4mg/mL, 1% SDS, 10 mM Tris, 10 mM EDTA, 5mM NaCl) and purified using QIAgen DNEasy® Plant Mini kit columns following an established protocol (Vallebueno-Estrada *et al*., 2023; Swarts *et al*., 2017).

Uracil-DNA-glycosylase (UDG) libraries were constructed at the ancient DNA facilities of Vienna University, Austria from 20 µL of ancient maize DNA following a published protocol tested in maize (https://github.com/Vallebueno/AncientLabProtocols.git) (Vallebueno-Estrada *et al*., 2023, Kentaro *et al*., 2013) with modifications as suggested in (Meyer *et al*., 2012). Libraries were amplified for 10 cycles with unique combinations of two indexing primers (Kircher *et al*., 2012). The quality of libraries was tested using Qubit 2.0 fluorometer (Thermo Fisher) and a High Sensitivity DNA Assay Chip Kit (Agilent, Waldborn Germany) on a Bioanalyzer 2100 (Agilent Technologies). UDG double index DNA Illumina libraries were sequenced using Novaseq S4 PE100 at Vienna Biocenter Facilities, Austria.

### Read processing, mapping, and genotyping

Double Index sequences of 8 nucleotides were used to tag libraries described above. Only reads with the correct index combination were used in downstream analysis. All libraries were filtered to remove adaptors and low-quality reads using Cutadapt (V1.13) (Marcel *et al*., 2011) and keeping reads longer than 30 bp with a quality above 10 Phred score. To remove possible molecular damage present in the UDG corrected libraries 2 bp of each side of the reads were trimmed using Cutadapt (V1.13) (Marcel *et al*., 2011). All samples were mapped to the current maize reference genome B73_V5 (Hufford *et al*., 2021) using BWA-MEM v 0.7.17 (Li *et al*., 2010).

Reads with multiple hits were removed using SAMtools map quality filters. As a clonal removal strategy, sequence duplication in reads was filtered with the rmdup function of SAMtools (V1.5) (Li *et al*., 2009). To eliminate reads with low certainty assignments, reads with a lower Phred score than 10 were removed using SAMtools (V1.5) (Li *et al*., 2009). Variation information was extracted and called using the mpileup and bcftools functions of SAMtools (Li *et al*., 2009). Genotype variants were then production called from the mpileup files based on previously observed SNPs using a custom HapMap build (Vallebueno-Estrada *et al*., 2024) comparable to present in previously published HapMap panels (Chia *et al*., 2012; Grzybowski *et al*., 2023; Bukowski *et al*., 2018).

### Gene diversity of ancient samples at genes with root phenotype

Based on the list of genes and coordinates of genes related to root phenotype (Data S8) allelic variant frequencies were calculated per site for each group of *Zea mays ssp. parviglumis* (n=70), *Zea mays ssp. mexicana* (n=81), *Zea mays ssp. mays* landraces (LR, n=94), and *Zea mays ssp. mays* ancient samples from different sites of Tehuacan valley (Coxcatlán cave, El Riego cave, San Marcos cave) from different ages (5,500 [n=7], 5,000 [n=2], 4,500 [n=1], 4,000 [n=2], 3,500 [n=3], 2,500 [n=6], 2,000 [n=6], 1,500 [n=5], 1,000 [n=2], 500 [n=9] calibrated years before present). All samples have been dated directly using accelerator mass spectrometry (Data S7). Dates were calibrated using OxCal v 4.4.4 (Ramsey *et al*., 1995), and corroborated with CALIB v 8.2 (Stuiver *et al*., 1993), to adjust the radiocarbon determinations to the IntCal20 Northern Hemisphere Radiocarbon Age Calibration Curve (Reimer *et al*., 2020). This adjustment was necessary to correct the dates and estimated age ranges based on the fluctuating atmospheric C14 concentrations. Delta C13 reflects that the material dated comes from a C4 plant as PEP carboxylase has a higher affinity for 13C isotope than Rubisco, thus the C14 determination likely comes from maize. Modern landrace samples (Data S9) were obtained from published Bioprojects: PRJNA300309, PRJNA479960, PRJNA641489 and HapMap 2 (Chia *et al*., 2012). Allelic frequencies were normalized per group and site based on the number of individuals within a group with non-missing information at a given position.

### Model limitations

Our approach presents several limitations, for instance, ancient phenotypes as well as ancient DNA might have been affected through time, and few ancient samples exist. Functional-structural models are a simplification of reality and cannot include all factors. For instance, teosinte in nature can grow in competition with other wild plants and can be subject to biotic stresses that are not considered in our model, nor are microbial symbionts such as mycorrhizas. We used extant teosintes to parameterize the model since ancient teosinte samples do not exist. Direct phenotypic plasticity (i.e. programmed growth responses to environmental stimuli rather than indirect plasticity from resource limitation (Gaudin *et al*., 2011; Hwang *et al.,* 2024) is not considered in our simulations. Soil O_2_ and soil temperature are not considered. Tillering was not simulated for teosinte. Simulations are run for 40 days and do not consider flowering time. Increase of nitrogen levels due to pre-Columbian fertilization was not considered due to a lack of archaeological evidence. Due to the nature of the analysis, it is impossible to measure fitness as reproductive success, instead, we are considering access to limiting resources (C, N, water) as a measure of fitness (Allaby *et al*., 2022). While simplification of the parameter space is an inevitable aspect of heuristic modeling, despite such simplifications, in silico results from *OpenSimRoot* and its predecessor *SimRoot* have shown excellent agreement with empirical results in previous studies (Lynch *et al*., 2023).

## Results

### Kernel weight increased by 6,000 yBP

Abundant seed reserves permit the development of increased SRN in maize and other domesticated grasses (Perkins and Lynch, 2021). Teosinte has an average kernel weight of 0.05 g, limiting the development of seminal roots. To understand when the kernel weight was sufficient to sustain increased SRN as kernel weight increased under domestication, we constructed a regression model with kernel weight as the response variable and width and length of the glume as predictors and used it to predict the kernel weight of several ancient specimens. We predicted that kernel weight of all ancient groups was greater than 0.2 ± 0.029 g, therefore suggesting that kernel weight was not a restriction to produce an increased seminal root number for ancient specimens since 6,000 yBP (Figure 2B). To test model accuracy, we predicted the seed weight of 31 individuals from a Multiparent Advanced Generation Inter-Cross maize population (Figure 2A). We obtained a significant positive correlation between the real and predicted kernel weight (R^2^=0.23, p = 0.0067). This indicates that the kernel weight of maize 6000 yBP was similar to modern maize, suggesting that 6000 yBP kernel weight was not a limitation to produce an increased SRN.

**Figure 2.**
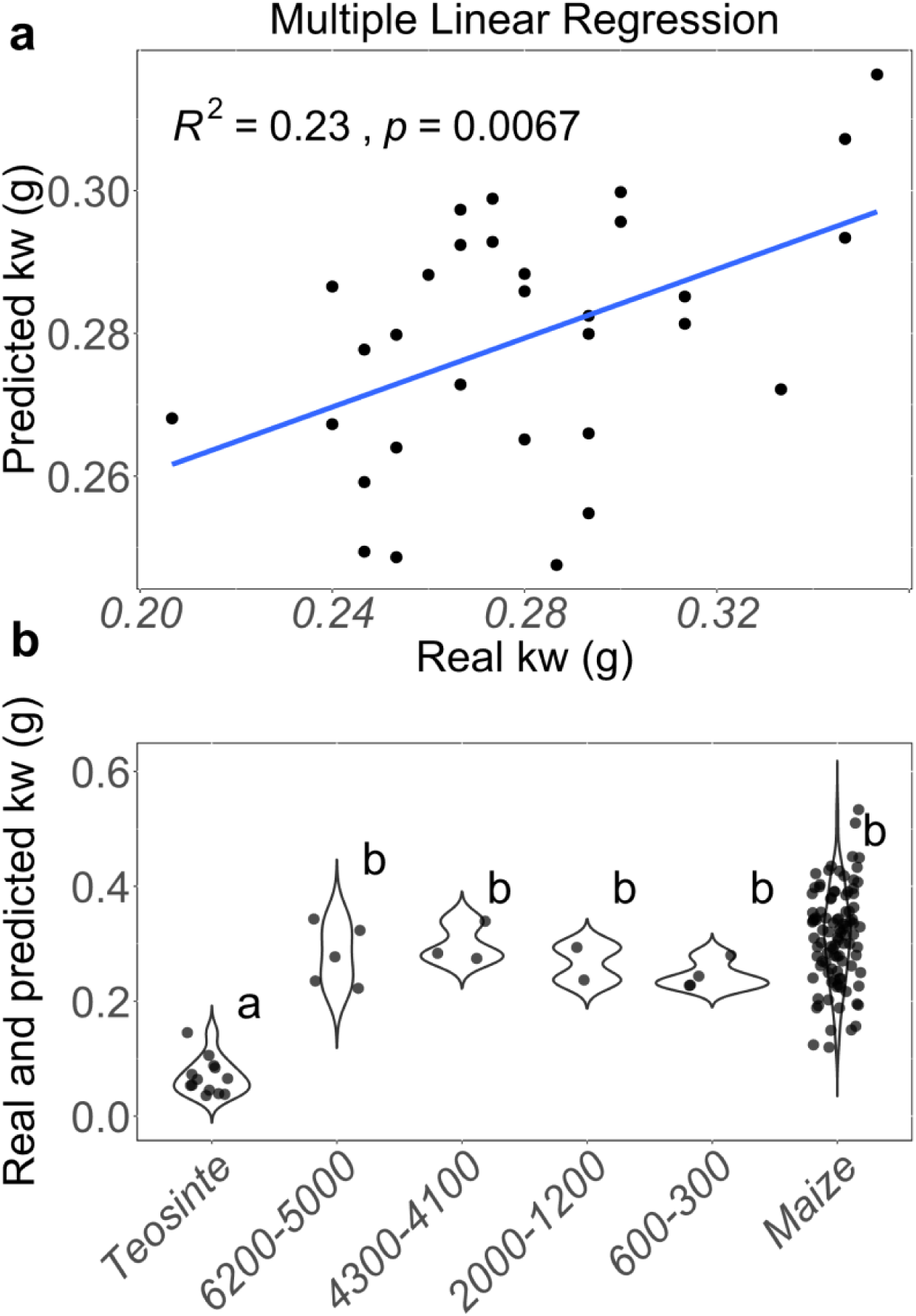
Prediction of kernel weight in ancient maize specimens from Tehuacan-Oaxaca dating from 6500 to 300 yBP. (a) Validation of multiple linear regression model constructed to predict kernel weight using glume width and length as predictors. (b) Comparison of predictions of kernel weight of ancient cobs dating from 6500 to 300 yBP with real kernel weight of teosinte and maize (Torres-Rodríguez *et al*., 2018; Ramos- Madrigal *et al*., 2016; Piperno and Flanery, 2001). Ancient samples were grouped based on their antiquity in 4 groups with ranging from 6200 to 5000, 4300 to 4100, 2000 to 1200 and 600 to 300 yBP. ANOVA and Tukey tests were performed to evaluate differences among groups. Significant differences in means between violin plots are represented as different letters with P ≤ 0.05.

### Seminal root number increased around 3,500 yBP

Ancient roots from the Tehuacan Valley showed that seminal roots were absent 5,000 yBP, suggesting the late fixation of this phenotype (Lopez-Valdivia *et al*., 2022). Our results showed that kernel weight was large enough and did not limit the production of seminal roots 6,000 yBP. However, it still remains unclear when the SRN increased. To determine when SRN increased in the last 6,000 years, we compared the allelic frequency of genes associated with SRN (Zm00001d021572, BIGE1 [BIG EMBRYO 1], and RUM1 [ROOTLESS WITH UNDETECTABLE MERISTEMS 1] [Wang *et al*., 2023; Suzuki *et al*., 2015; Salvi *et al*., 2016; Woll *et al*., 2005; Suzuki *et al*., 2015]) among populations of *teosinte parviglumis*, *teosinte mexicana*, modern landraces and archeological maize samples from 5,500, 5,000, 4,500, 4,000, 3,500, 2,500, 2,000, 1,500, 1,000, and 500 yBP.

We found a total 84, 172, and 165 SNPs for Zm00001d021572, BIGE1, and RUM1 in the ancient samples (Data S10). To obtain a reference of the SNPs that changed trough domestication we filtered the SNPs that had at least 40% difference in the allelic frequency when comparing between maize and teosinte populations. We obtained 14, 4, and 1 SNPs that showed a shift from teosinte to maize for Zm00001d021572, BIGE1, and RUM1, respectively (Figure S1-S3). For Zm00001d021572, 8 of 14 SNPs showed an allelic frequency similar to teosinte in ancient samples corresponding to 5,500 yBP, but then switched to an allelic frequency similar to maize in samples from 3,500-500 yBP, while the other 6 SNPs showed a modern allele in all ancient samples (Figure S1). BIGE1 showed that 3 of 4 SNPs had a teosinte-like frequency in samples from 5,000 yBP, and maize-like allelic frequency in samples from 3,500-500 yBP (Figure S2). RUM1 only showed 1 SNP which was different in maize and teosinte populations and the ancient samples showed the modern allele (Figure S3). While direct genotype-phenotype correlations remain elusive, the transition from teosinte-like to maize-like allelic frequencies in Zm00001d021572 and BIGE1 between 5,500 to 3,500 yBP suggests that the SRN might have increased between 5,500 to 3,500 yBP.

### Reduced NRN and MCS appeared around 8,000 yBP

Tehuacan Valley maize roots showed reduced NRN and MCS by 5,000 yBP (Lopez-Valdivia *et al*., 2022). However, the temporal order of appearance of reduced NRN and MCS between 10,000 to 5,000 yBP is unknown. To find the most parsimonious scenario of the appearance of reduced NRN and MCS, we used *OpenSimRoot* to simulate phenotypic combinations of MCS and NRN in the environments corresponding to 8,000 and 6,000 yBP (Figure 3A). The scenario in which reduced NRN and MCS appeared by 8,000 yBP (scenario C) has the greatest shoot dry weight production compared with teosinte (Figure 3B). *OpenSimRoot* is a heuristic tool designed to assess the potential effects of root phenotypes on soil resource capture. If we assume that greater ability to acquire limiting resources is associated with fitness (Allaby *et al*., 2022), we can suggest that scenario C will have the maximum integral biomass over time. The appearance of reduced NRN by 8,000 yBP and then MCS by 6,000 yBP (scenario A) is also possible because it produces the second-greatest shoot dry weight compared with teosinte (Figure 3). The scenario where reduced NRN and MCS appeared by 6,000 yBP (scenario D) produced a similar shoot biomass to teosinte, being the third most probable scenario. It is unlikely that MCS appeared earlier than reduced NRN because the shoot biomass produced by scenario B is less than teosinte, giving no advantage in the environment of 8,000 yBP (Figure 3B). The appearance of reduced NRN and MCS by 8,000 yBP is the most likely scenario; however, we cannot discard the possibility that reduced NRN appeared earlier than MCS. Finally, the poor fitness of scenario B suggests it is unlikely that MCS appeared earlier than reduced NRN.

**Figure 3.**
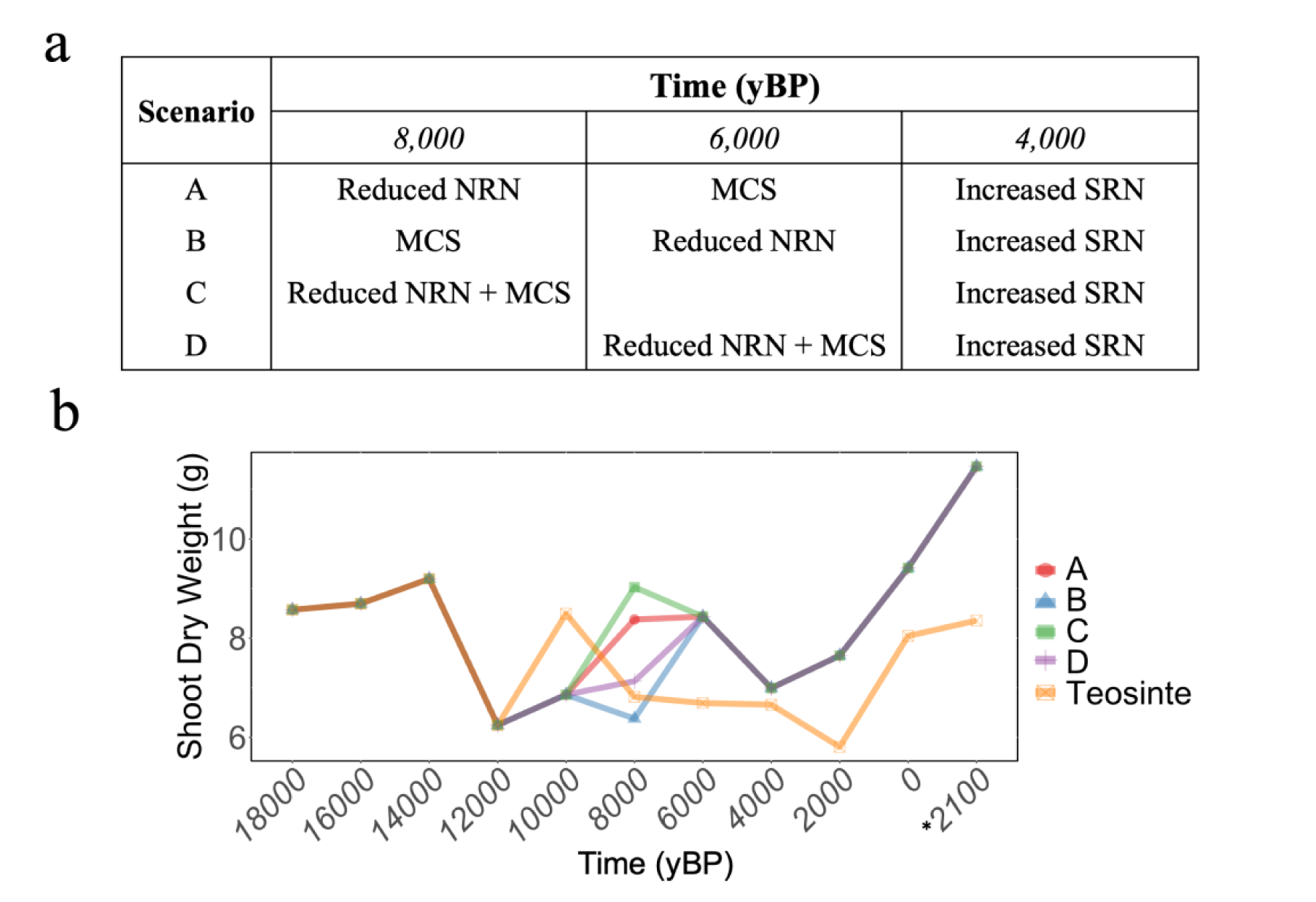
Scenarios for the root phenotypic transition from teosinte to maize. (A) Scenarios correspond to combinations of reduced NRN, MCS, and increased SRN through time. (B) Simulated shoot dry weight produced by scenarios A, B, C, D and teosinte in the environments corresponding to the last 18,000 yBP. The future scenario denoted by * corresponds to 2100 common era and was simulated based on climatic projections (61).

### Nitrogen availability shifted to deeper soil domains with the emergence of irrigation

Archeological evidence shows that irrigation systems were implemented in the Tehuacan Valley by 6,000 yBP (Neely *et al*., 2022). To understand the effect of irrigation on the nitrogen dynamics in Tehuacan soils during the last 18,000 years, we simulated the soil in 2,000-year timesteps with *OpenSimRoot* considering the implementation of irrigation and soil degradation. We found that nitrate distribution increased its depth from 0-20 cm to 30- 50 cm after the implementation of irrigation systems 6,000 yBP (Figure 4A). When we simulated soil degradation by reducing epipedon depth and organic matter content (McAuliffe *et al*., 2021; Araus *et al*., 2014; Cook, 1949) after 6000 yBP we observed that nitrate concentration decreased from 8 mM to 1.5 mM. We also observed that during simulated drought periods (16,000, 12,000, and 8,000 yBP) the soil domains with greater nitrate concentration tend to be distributed between 0-10 cm, while in wet periods (14,000, 10,000 yBP) nitrate-rich regions are distributed between 10-20 cm. We also observed that during simulated drought periods, root systems experienced greater penetration resistance as they encountered dryer, harder soil domains compared with wet periods (Figure 4A). Interestingly, we observed that soil domains with the greatest nitrogen availability switched from 40-60 cm in modern environments to 0-20 cm in projected future environments. In contrast, water availability, represented as the hydraulic head, is reduced by 200% and located 40 cm deeper in future environments. Irrigation and fertilization in modern high-input agriculture increased nitrogen concentration but shifted the nitrogen availability to deeper soil domains.

**Figure 4.**
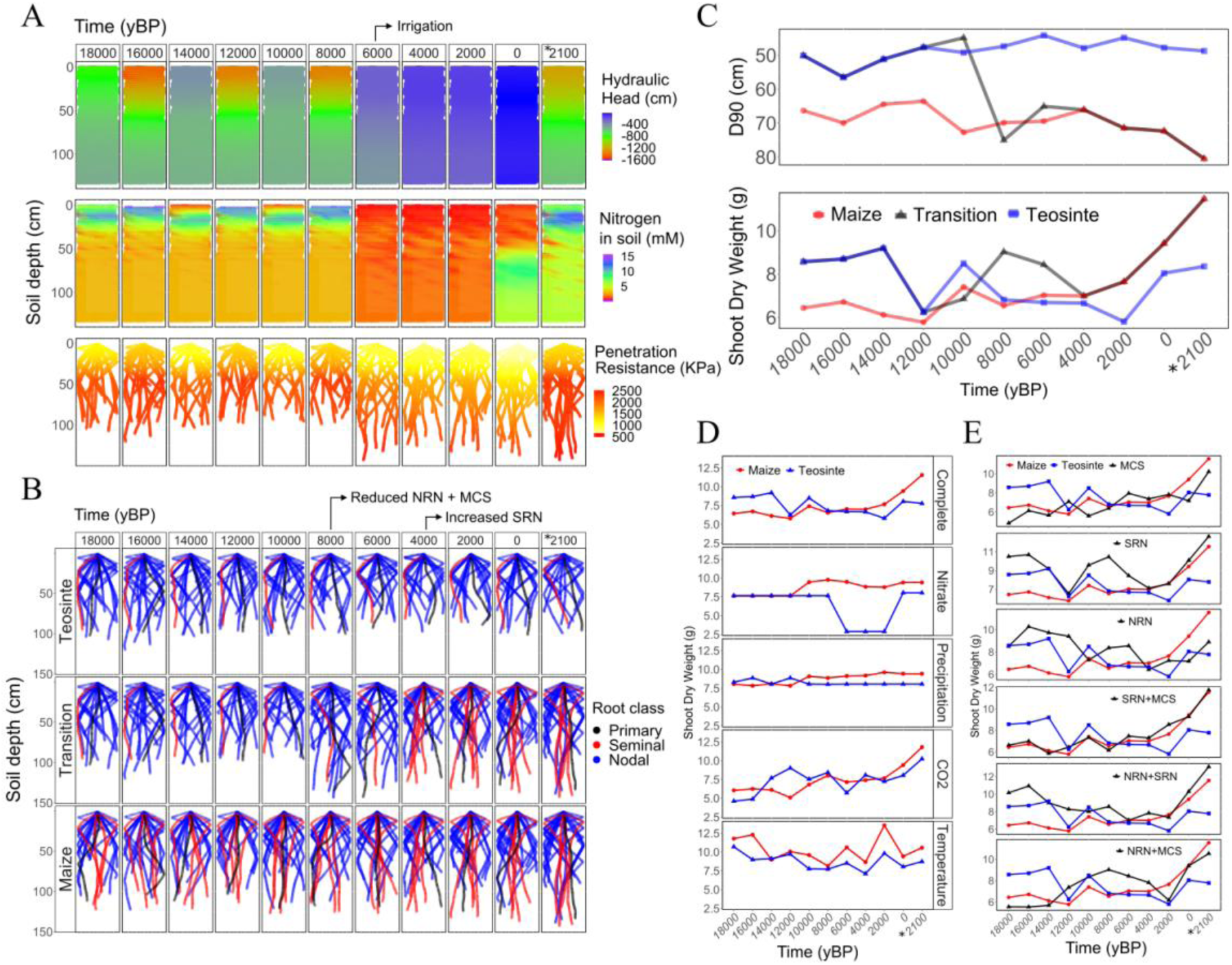
Simulations of root phenotypes and soil environments in the last 18000 years BP. (a) Soil environment matrix during the last 18,000 yBP represented as the hydraulic head (cm), nitrogen concentration (mM), and penetration resistance (kPa) at day 40. (b) Root phenotypes of teosinte, transitional phenotype (Scenario C), and maize simulated over the environments corresponding to the last 18,000 yBP. (c) Simulated D90 [soil depth at which 90% of the root length was above] and shoot dry weight of root phenotypes of teosinte, transitional phenotypes and maize in the environments of the last 18,000 yBP. (d) Environmental dissection into nitrogen availability, precipitation, atmospheric CO_2_, and temperature. (e) Phenotypic dissection of MCS, SRN, and NRN and their interactions over time. The future scenario denoted by * corresponds to 2100 common era and was simulated based on climatic projections (Lee *et al*., 2021).

### Root phenotypes adapted to irrigation

We present a teosinte-to-maize root phenotypic transition, which condenses our previous results where reduced NRN and MCS likely appeared by 8,000 yBP, kernel weight was as large as maize by 6,000 yBP, and increased SRN appeared by 3,500 yBP (Figure 4B). To evaluate the role of the root phenotypic transition in the last 18,000 yBP, we simulated the interaction of the roots of teosinte, maize, and transitional phenotypes with environments of that period. Teosinte produced greater shoot biomass than maize between 18,000 to 10,000 yBP, while maize produced greater shoot biomass than teosinte from 8,000 yBP to present (Figure 4C). Transitional phenotypes had greater shoot biomass than maize in the late Pleistocene and greater shoot biomass than teosinte in Holocene environments. In general, the D_90_ [soil depth attained by 90% of root length] was 50-60 for teosinte and 65- 80 cm for maize. Interestingly, the D_90_ of the transitional phenotype was 50-60 cm between 18,000 to 10,000 yBP and then increased to 65-80 cm after 8,000 yBP, following the distribution of nitrate over time. Maize outperformed teosinte in modern environments both with and without irrigation (Figure S4). In a simulated future environment, the maize phenotype produced 30 cm deeper roots, matching with the availability of water, and produced 28% greater biomass compared with the teosinte phenotype. The transitional phenotype increased root depth after 8,000 yBP, matching the deeper distribution of nitrate after the advent of irrigation.

### Phenotypic dissection suggests an ordered appearance of root phenotypes

The teosinte-to-maize root phenotypic transition includes reduced NRN, MCS, and increased SRN, interacting with varying nitrogen availability, atmospheric CO_2_ concentration, temperature, water availability (both precipitation and irrigation), and soil mechanical impedance. To understand phenotype-environment interactions during the transition of root phenotypes, we dissected root phenotypes and environments and simulated their individual and combined effects over 18,000 years. Our simulations showed that nitrogen had the greatest impact on the performance of maize over teosinte when compared with CO_2_, temperature, and precipitation/irrigation (Figure 4D). Interestingly, the adaptive advantage of MCS increased linearly with atmospheric CO_2_ concentration (Figure 4E). The combination of MCS with reduced NRN improved plant biomass as early as 12,000 yBP (Figure 4E). While reduced NRN alone improved biomass during the late Pleistocene and early Holocene, this benefit decreased by the late Holocene. However, reduced NRN combined with increased SRN recovered utility by the late Holocene (Figure 4E), suggesting a crucial role of increased SRN in adapting to the conditions of the late Holocene. Intriguingly, increased SRN alone showed utility as early as 18,000 yBP. This finding contrasts with the absence of evidence for increased SRN in 5,000 yBP ancient roots and the shift in allelic diversity of genes associated with seminal root number around 3500 yBP. This discrepancy highlights the potential lag between the theoretical advantage of these phenotypes and its actual appearance in the archaeological record. In summary, our findings indicate that nitrogen was the most important environmental factor that limited the utility of modern maize root systems in the Holocene in Tehuacan Valley, while an ordered appearance of reduced NRN, MCS, and increased SRN was necessary for the adaptation of plant populations to the modern agricultural environment.

## Discussion

We propose a model for the gradual transition of root phenotypes from teosinte to maize. First, reduced NRN and MCS appeared between 12,000 to 8,000 yBP. Then, kernel weight was similar to modern maize by around 6,000 yBP, and finally, increased SRN likely appeared at around 3,500 yBP. This gradual transition reflects a dynamic interplay between root phenotypes and the environment, especially nitrogen. Nitrogen-rich soil domains shifted from shallow to deeper depths after the advent of irrigation. In turn, maize phenotypes switched from a shallow to a deeper distribution of root length during domestication, matching the distribution of soil nitrogen. Finally, an ordered appearance of reduced NRN, MCS, and increased SRN was necessary for adaptation to Holocene agriculture in the Tehuacan Valley, where nitrogen distribution was the main environmental factor influencing adaptation.

We used functional-structural modeling to evaluate our hypotheses, because of the inaccessibility of this topic to empirical research. Empirical evidence supports the environmental transitions associated with maize domestication we model here of changing precipitation, temperature, atmospheric CO_2_, soil degradation, nitrogen bioavailability, tillage, and irrigation. However, it would not be feasible to recreate the array of environments needed to empirically dissect the isolated and interacting effects of these factors on plant fitness. Likewise, empirical data from ancient maize root specimens from the Tehuacan Valley is used to parameterize our models, however, it would not be feasible to recreate an array of root phenotypes varying in specific ways to empirically test the effects of these phenotypic transitions alone and in combination on plant fitness in contrasting environments. Furthermore, a very limited number of ancient root specimens exist that do not capture the entire domestication sequence. Analysis of a multidimensional decision space of phenotypes that do not currently exist (including hypothetical phenotypes that may never have existed) interacting with environments that no longer exist (and may never have existed, such as ancient soils with modern atmospheric CO_2_) is ideally suited to *in silico* approaches. The tool we use is *OpenSimRoot*, a feature-rich, heuristic, functional-structural plant-soil model which has proved useful in understanding several aspects of root/soil interactions (*e.g.* Ajmera et al., 2022; Strock et al., 2022; Schafer et al., 2022; Lopez-Valdivia et al., 2023, Rangarajan and Lynch 2024) and the root phenome (*e.g.* Rangarajan and Lynch, 2021; Nielsen et al., 1997). Heuristic models are useful primarily for the insights they provide rather than for their ability to predict empirical outcomes (Wullschleger et al., 1994; Yin and Struik, 2008; Zhu et al., 2016), emphasizing the validity of underlying processes and parameters rather than agreement with empirical measurements (Dunbabin et al., 2013). The suitability of *OpenSimRoot* for this study lies in its ability to simulate the interaction of root growth and soil processes dynamically in time and space while considering the effects of plant resource availability and allocation and its link with water and nutrient capture. Specifically, *OpenSimRoot* explicitly models the interacting effects of the key variables relevant to our hypotheses, including the effects of atmospheric variables (*i.e.*, temperature, humidity, CO_2_ concentration, light intensity and duration, and precipitation) on plant and soil processes, the effects of the soil environment (*i.e.*, soil texture, water content, mechanical impedance, organic matter content, and the transport and bioavailability of nitrogen, phosphorus, and water) on root growth and soil resource capture, and the effects of root architectural and anatomical phenotypes on soil resource capture in response to the soil environment. Our results support the hypothesis that the evolution of maize root phenotypes was driven by anthropogenic changes in the soil environment, propose a sequence of evolutionary adaptations in root phenotypes that may have optimized plant fitness, and indicate that nitrogen availability was a key environmental factor shaping maize root evolution. These results provide a conceptual framework for understanding root evolution during maize domestication, and generate specific hypotheses that merit empirical validation.

Maize seedlings require at least 0.07 g of seed carbohydrate reserves to sustain increased SRN (Perkins and Lynch, 2021). The limited seed carbohydrate reserves of teosinte (Flint-Garcia, 2017; Flint-Garcia *et al*., 2009) presumably limit the production of seminal roots. Interestingly, greater SRN is associated with nitrogen and phosphorus capture in maize seedlings (Perkins and Lynch, 2021; Flint-Garcia *et al*., 2009) and drought stress tolerance (Yu *et al*. 2024). Our results suggest that the kernel weight of early domesticates was as large as modern maize by 6,000 yBP, matching temporally with the teosinte *mexicana* introgression associated with modern maize kernel phenotypes (Yang *et al*., 2023). This finding raises the question of: why increased SRN was absent at 5,000 yBP, even though seed reserves were sufficient? Our results show that the allelic frequencies of genes associated with SRN were similar to teosinte in ancient samples from 5,000 yBP but shifted to maize-like frequencies in samples between 3,500-500 yBP. This suggests that SRN increased around 3,500 yBP, potentially driven by increasing population pressure and the intensification of agriculture in the Tehuacan Valley. The human population of the Tehuacan Valley increased from 2,000 to 80,000 between 3,500-1,300 yBP accompanied by an increased dependence of agricultural products in their diets (MacNeish, 1967). Intensive irrigated agriculture (Neely et al., 2022) and consequent soil degradation (McAuliffe et al., 2001) might have increased the selection pressure for root phenotypes that improved adaptation to nitrogen stress. While our study focuses on the Tehuacan Valley, similar population increases occurred in regions of Belize and Guatemala, where increased maize consumption was likely driven by dry seasons during the late Holocene (Ray et al., 2023). This suggests that the intensification of agriculture and the associated selection pressures on maize roots may have been a widespread phenomenon. In summary, our simulations show that the appearance of increased SRN by 3,500 yBP would have improved plant growth in degraded soil with irrigation, allowing increased crop production and the consequent demographic explosion.

Yu *et al*. (2024) proposed that seed size is independent of seminal root number (SRN) in maize, based on evidence of traditional maize varieties and *su1* and *sh2* mutants, showing no correlation between seed size and SRN. However, teosintes were not included in this comparison. This is relevant because 95% of maize accessions have larger seed reserves than most teosintes (Figure S5). The smallest seed reserves of maize landraces have the potential to develop at least 4 seminal root number (Perkins and Lynch, 2021), the average seminal root number reported for maize (Burton et al., 2013). Although a direct association between seed size and increased SRN in maize remains unclear, the potential influence of seed size on SRN, particularly during the evolutionary transition from teosinte to maize, requires further investigation.

The gene *Teosinte Branched 1* (*Tb1*), associated with NRN (Gaudin *et al*., 2014), presented the allelic frequency of modern maize by 5,000 yBP (Vallebueno-Estrada *et al*., 2016), aligning with the reduced NRN phenotype observed in contemporary specimens (Lopez-Valdivia *et al*., 2022) and with the non-tiller ancient roots found in Tehuacan (MacNeish, 1967). Increased NRN represents a metabolic cost because roots are heterotrophic (Lynch, 2013). However, costly root phenotypes are selected in a competitive environment if they confer greater access to limited resources (Allaby *et al*., 2022, Henry *et al*., 2010; Zhang *et al*., 2014; Postma and Lynch, 2012; Rubio *et al*., 2001; Rubio *et al*., 2003). For instance, maize genotypes with increased NRN have greater biomass under phosphorus stress (Sun *et al*., 2018), while maize genotypes with less NRN produced greater biomass under nitrogen (Saengwilai *et al*., 2014) and drought stress (Gao *et al*., 2016). The contrasting utility of NRN hinges on the spatiotemporal coincidence between root foraging and resource availability in the soil. Plants with reduced NRN invest resources in making fewer but longer nodal root axes, which can reach deeper soil domains and capture leached nitrogen and water (Saengwilai *et al*., 2014; Gao *et al*., 2016). In contrast, phosphorus is an immobile resource and has greater bioavailability in the topsoil. Therefore, increased NRN increases root length in the topsoil and improves phosphorus capture (Sun *et al*., 2018). Reduced NRN reduces intraspecific competition and increases biomass in high-density environments (York *et al*., 2015), which would be desirable in ancient polycultures with bean and squash (Zhang *et al*., 2014; Postma and Lynch 2010) as prevalent in the Tehuacan Valley (MacNeish, 1967).

MCS was present in ancient roots from the Tehuacan Valley 5,000 yBP (Lopez-Valdivia *et al*., 2022). Our simulations show that the greater carbon cost of MCS formation becomes more burdensome with low atmospheric CO_2_ concentrations in the late Pleistocene and early Holocene. With the advent of irrigation about 6,000 yBP, water deficit stress was relieved at the cost of the loss of soil organic matter and greater nitrate leaching. In these conditions, MCS becomes advantageous as it permits axial roots to penetrate deeper, harder soil (Schneider *et al*., 2021), as would reduced NRN, which increases rooting depth and nitrate capture (Saengwilai *et al*., 2014). It is unlikely that MCS preceded reduced NRN because MCS combined with increased NRN becomes too metabolically costly. In contrast, if reduced NRN preceded MCS, the metabolic cost of MCS would be compensated by the reduced NRN. Reduced NRN combined with MCS was useful since 12,000 yBP (Figure 4E), suggesting they might be the earliest root phenotypes in maize domestication. Surprisingly, MCS was absent in *Z. mays ssp. parviglumis* and *Z. mays ssp. mexicana* (Schneider *et al*., 2021), possibly because of a role in interplant competition for topsoil resources in native soil, thereby favoring increased NRN without MCS. Overall, our results show how adaptation to a particular environment involves the interaction of several phenotypes (Schäfer *et al*., 2022; Ajmera *et al*., 2022; Lynch *et al*., 2023).

In summary, the evolution of root phenotypes is fundamental to understanding crop domestication. We provide support for the hypothesis that anthropogenic modification of the soil environment interacted with anthropogenic selection to shape the evolution of root phenotypes during maize domestication. Our results indicate that the appearance of root phenotypes was important for the adaptation of crops to cultivated environments in Holocene agriculture and provide guidance for the use of root phenotypes as breeding targets for future environments.

## Acknowledgments

The authors are grateful to the archeologists and sample curators at the Instituto Nacional de Antropología e Historia for facilitating access to ancient maize samples. We are also grateful to the High-Performance Computing Center as well as the IT service at Pennsylvania State University for facilitating the computational resources and technical support required for this investigation. We thank Kathleen Brown and Jesse Lasky for reviewing our manuscript and providing useful comments and suggestions. We also thank the Zeavolution online seminar for the insightful discussion and comments on this research. This project has received funding from the Foundation for Food and Agriculture Research under award number 602757 (ILV, HR, JSS, JPL), the United States Department of Agriculture National Institute of Food and Agriculture Federal Appropriations under Hatch Project PEN04732. (ILV, HR, JSS, JPL), the National Science Foundation award number 0091490 (BFB), the Social Sciences and Humanities Research Council of Canada (MB), and the European Union’s Framework Programme for Research and Innovation Horizon 2020 (2014-2020) under the Marie Curie Skłodowska Grant Agreement Nr. 847548 (MV, KS). We acknowledge that access to the Tehuacán collections for the radiocarbon dated ancient DNA analyses was granted by J. Garcia-Barcena, Presidente, Consejo de Arqueología, Instituto Nacional de Antropología e Historia in oficio 401-36/786 to Bruce Benz.

## Author Contributions

Conceptualization: JPL, ILV; methodology: ILV, HR, JSS, MVE, KS, BFB, MB, SPL, RJHS, HS, JPL; investigation: ILV, HR, JSS, MVE, HS, SPL, RJHS,

JPL; visualization: ILV, JPL; funding acquisition: JPL; project administration: JPL; supervision: JPL; writing – original draft: ILV, JPL; writing – review and editing: ILV, HR, JSS, MVE, KS, SPL, RJHS, HS, JPL.

## Data availability

All the input data required to replicate the simulations in *OpenSimRoot* is available in the following repository: https://github.com/ilovaldivia/Lopez-Valdivia_2023_RootEvolution.git and in supplementary materials.

## Supporting Information This section file includes

Figures S1 to S4 Legends for Movies S1

Legends for Datasets S1 to S9

## Other supporting materials for this manuscript include the following

Movies S1 Datasets S1 to S10

**Movie S1 (separate file).** Animation of Figure 1.

**Dataset S1 (separate file).** Glume width, glume length, and kernel weight for a portion of the MAGIC maize population.

**Dataset S2 (separate file).** Kernel weight predictions for ancient maize samples from Tehuacan Valley.

**Dataset S3 (separate file).** Raw atmospheric data for Tehuacan Valley (18°27′42″N, 97°23′34″W) corresponding to the precipitation, minimum temperature, maximum temperature, wind speed, and relative humidity from 1980 to 2021.

**Dataset S4 (separate file).** Average atmosphere patterns for a rainy season (June 19 – August 18) in Tehuacan Valley (18°27′42″N, 97°23′34″W) between 1980 and 2021. Data correspond to precipitation, minimum temperature, maximum temperature, wind speed, and relative humidity.

**Dataset S5 (separate file).** Soil texture and van Genuchten parameters from Tehuacan Valley. Core coordinates -97.3340, 18.3759, core number 215588, MX-INEGI dataset, SoilGrids (65).

**Dataset S6 (separate file).** Adjustments in soil and atmospheric parameters to simulate different eras based on paleoclimate and archeologic data.

**Dataset S7 (separate file).** Ancient samples and dating by accelerator mass spectrometry.

**Dataset S8 (separate file).** Root genes associated with seminal root number.

**Dataset S9 (separate file).** Maize samples genomic data

**Dataset S10 (separate file).** SNPs for genes Zm00001d021572, BIGE1, and RUM1 in the landraces (LR), teosintes (mexicana and parviglumis) and ancient samples of different ages (500 - 5500).

---

**Fig. S1.**
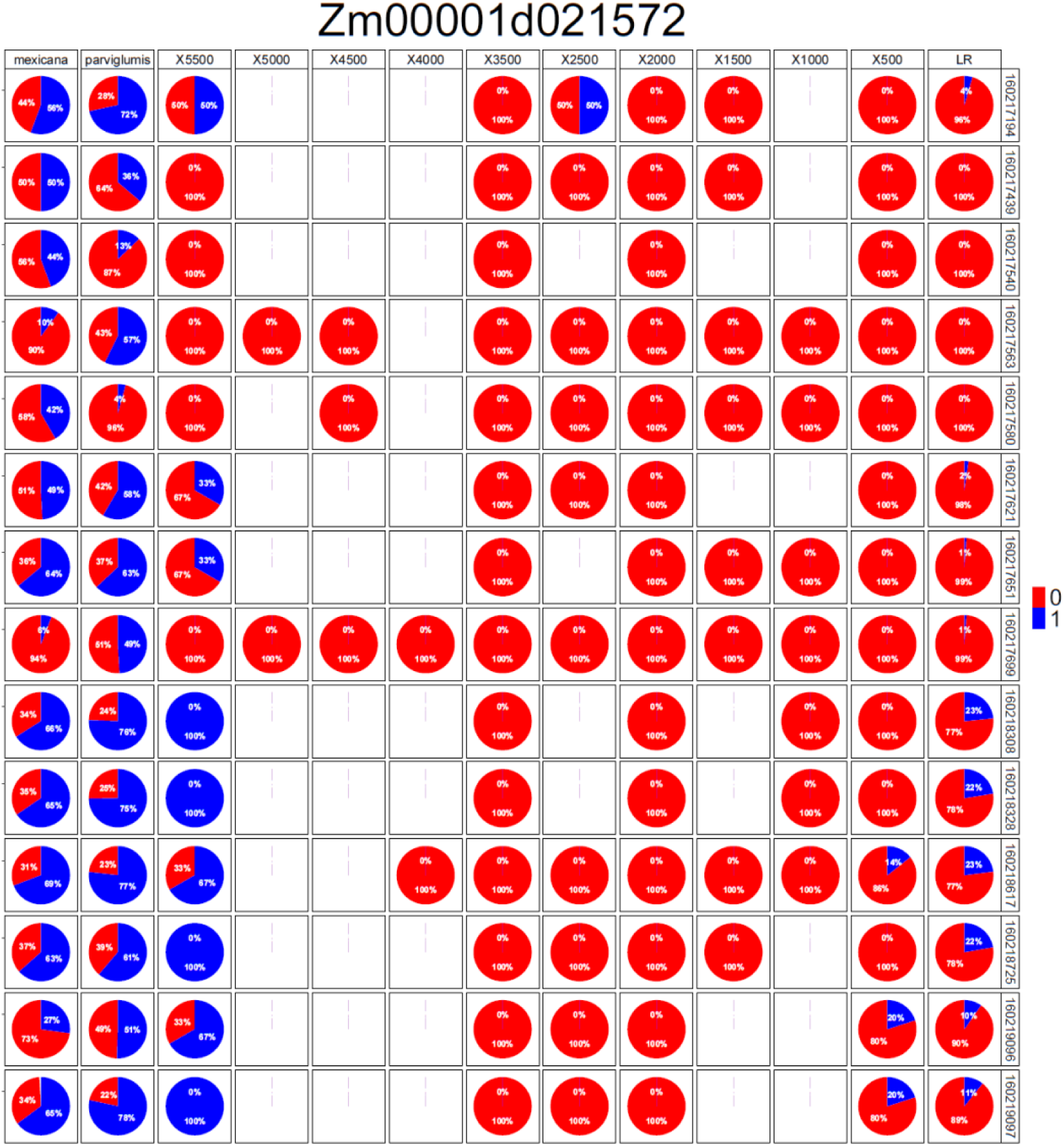
SNPs with at least a 40% difference in their allelic frequency when compared between maize and teosinte in the gene Zm00001d021572. Red represents the allele from the reference genome (B73) while blue represents an alternative allele. The absence of pie plots represents a lack of coverage in the ancient samples. Labels: mexicana = Zea mays ssp. mexicana (n=81), parviglumis = Zea mays ssp. parviglumis (n=70), LR = Zea mays ssp. mays (n=94), X5500 = ancient maize 5,585-4,975 yBP (n=7), X5000 = ancient maize 4,831-4,438 yBP (n=2), X4500 = ancient maize 4,871-3,581 yBP (n=1), X4000 = ancient maize 3,820-3,451 yBP (n=2), X3500 = ancient maize 4,849-2,324 yBP (n=3), X2500 = ancient maize 2,700-2,005 yBP (n=6), X2000 = ancient maize 2,051-1,410 yBP (n=6), X1500 = ancient maize 1,532-1,316 yBP (n=5), X1000 = ancient maize 676-560 yBP (n=2), X500 = ancient maize 529-157 yBP (n=9). All dates are expressed in calibrated yBP.

**Fig. S2.**
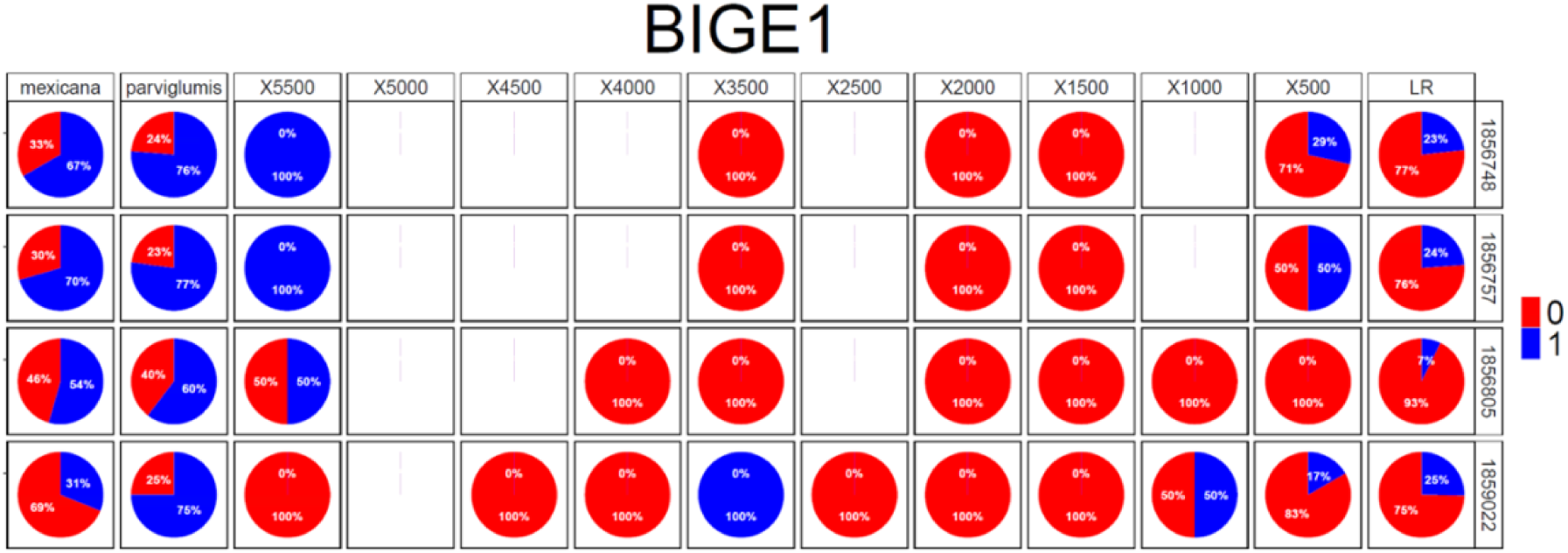
SNPs with at least a 40% difference in their allelic frequency when compared between maize and teosinte in the gene BIGE1. Red represents the allele from the reference genome (B73) while blue represents an alternative allele. The absence of pie plots represents a lack of coverage in the ancient samples. Labels: mexicana = Zea mays ssp. mexicana (n=81), parviglumis = Zea mays ssp. parviglumis (n=70), LR = Zea mays ssp. mays (n=94), X5500 = ancient maize 5,585-4,975 yBP (n=7), X5000 = ancient maize 4,831-4,438 yBP (n=2), X4500 = ancient maize 4,871-3,581 yBP (n=1), X4000 = ancient maize 3,820-3,451 yBP (n=2), X3500 = ancient maize 4,849-2,324 yBP (n=3), X2500 = ancient maize 2,700-2,005 yBP (n=6), X2000 = ancient maize 2,051-1,410 yBP (n=6), X1500 = ancient maize 1,532-1,316 yBP (n=5), X1000 = ancient maize 676-560 yBP (n=2), X500 = ancient maize 529-157 yBP (n=9). All dates are expressed in calibrated yBP.

**Fig. S3.**
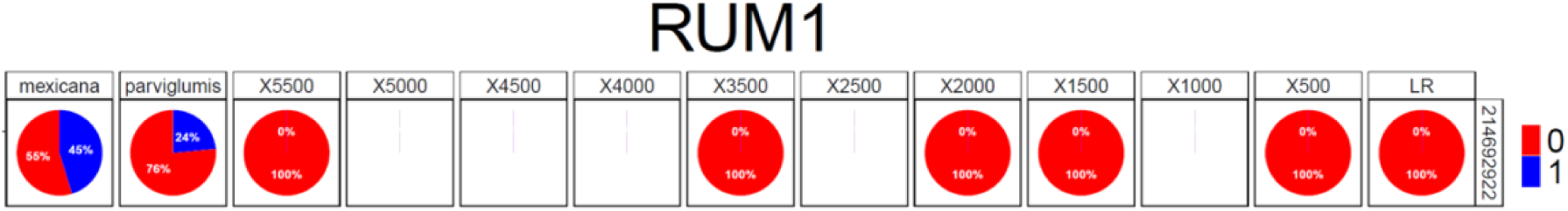
SNPs with at least a 40% difference in their allelic frequency when compared between maize and teosinte in the gene RUM1. Red represents the allele from the reference genome (B73) while blue represents an alternative allele. The absence of pie plots represents a lack of coverage in the ancient samples. Labels: mexicana = Zea mays ssp. mexicana (n=81), parviglumis = Zea mays ssp. parviglumis (n=70), LR = Zea mays ssp. mays (n=94), X5500 = ancient maize 5,585-4,975 yBP (n=7), X5000 = ancient maize 4,831-4,438 yBP (n=2), X4500 = ancient maize 4,871-3,581 yBP (n=1), X4000 = ancient maize 3,820-3,451 yBP (n=2), X3500 = ancient maize 4,849-2,324 yBP (n=3), X2500 = ancient maize 2,700-2,005 yBP (n=6), X2000 = ancient maize 2,051-1,410 yBP (n=6), X1500 = ancient maize 1,532-1,316 yBP (n=5), X1000 = ancient maize 676-560 yBP (n=2), X500 = ancient maize 529-157 yBP (n=9). All dates are expressed in calibrated yBP.

**Fig. S4.**
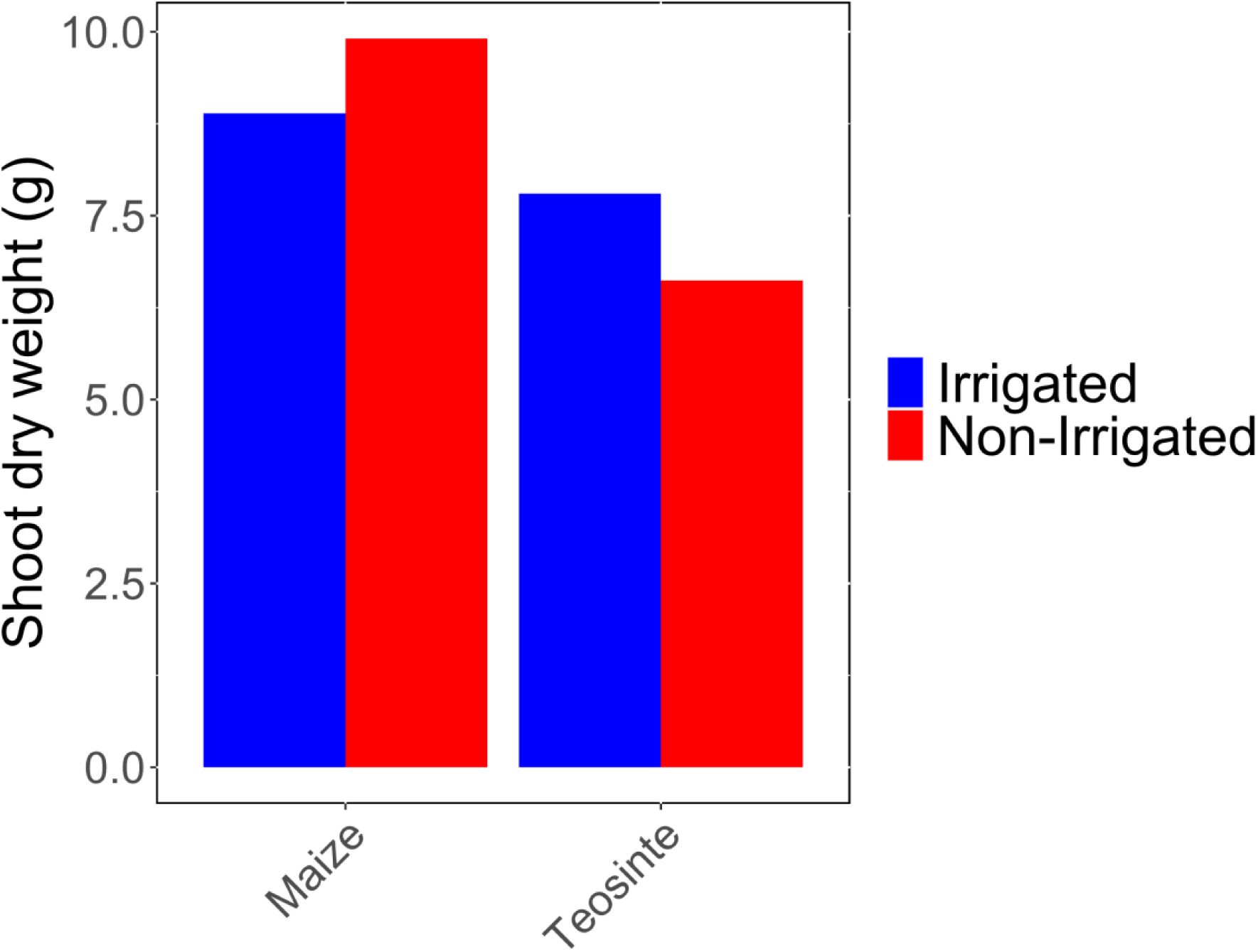
Simulation of the modern environment in Tehuacan Valley under irrigation and non-irrigation conditions.

**Fig. S5.**
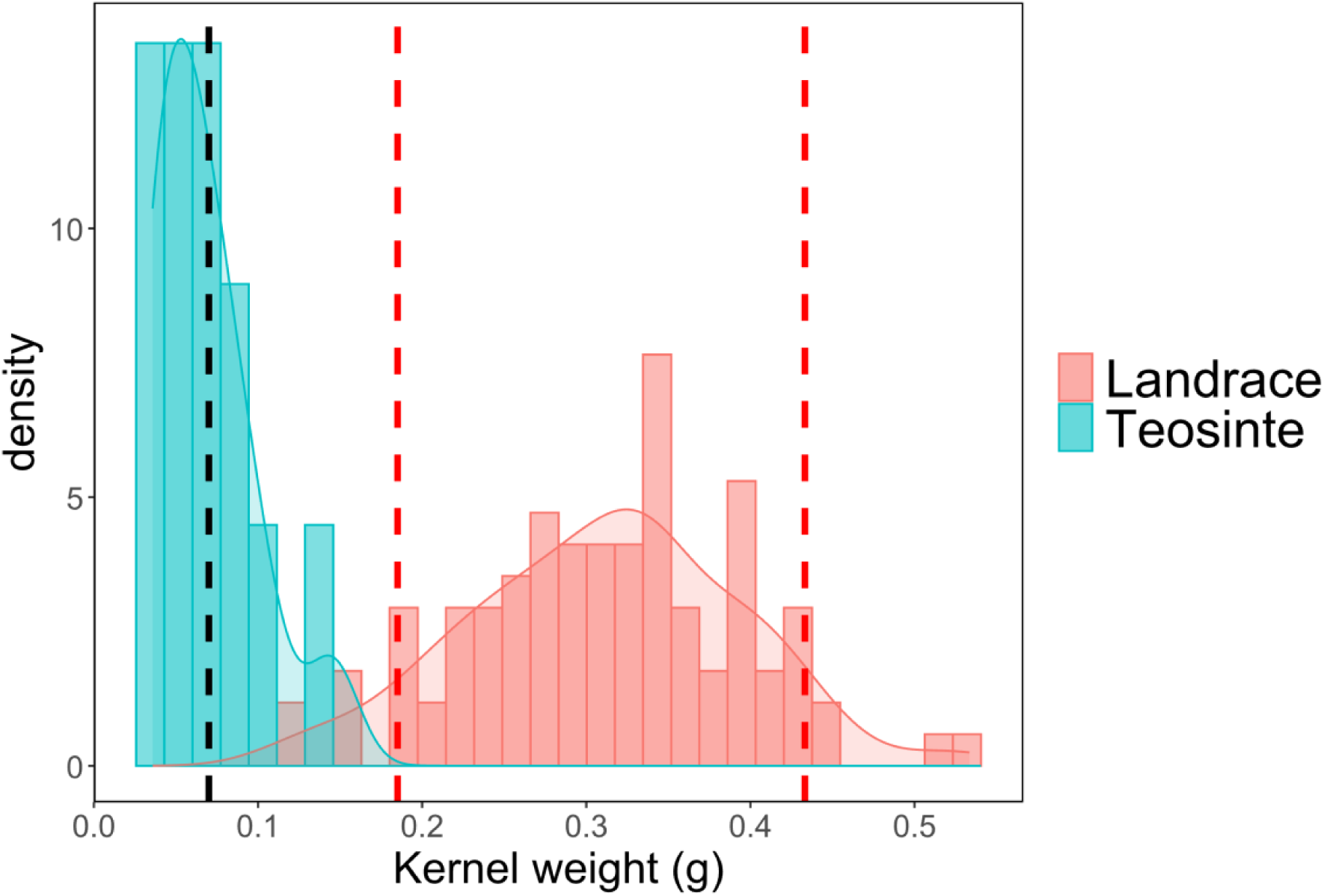
Distribution of kernel weight in 99 maize landraces and 13 teosintes. Kernel weight was determined as the average weight of five kernels per accession from Burton *et al*., 2013. Dashed red lines indicate the 5th and 95th percentiles of the kernel weight distribution in maize landraces. Black dashed line indicates the minimum kernel weight required to produce 4 seminal roots (Perkins and Lynch, 2022), reported as the average seminal root number for *Zea mays L. subsp. mays* (Burton *et al*., 2013).

